# Development of a Multifunctional Extracorporeal Life Support (ECLS) System for Lung and Kidney Support: The Pneuma-K ECLS System

**DOI:** 10.1101/2024.11.06.621276

**Authors:** Jian Kwon, Daniel J. Weiss, Patrick C. Lee, Jake K. Lee

## Abstract

Multi-organ failure (MOF), particularly in the coexistence of acute kidney injury (AKI) and acute lung injury (ALI), presents a significant challenge in intensive care units (ICU) and is associated with exceedingly high mortality rates. Respiratory and renal failures are frequently managed by extracorporeal membrane oxygenation (ECMO) and continuous renal replacement therapy (CRRT), respectively. However, employing these therapies using separate devices requires specialized facilities, adds to complexity, and increases the risks of clotting due to the extensive artificial surface areas involved. Therefore, an integrated device capable of providing simultaneous respiratory and renal support is essential. This paper introduces the Pneuma-K ECLS system, which incorporates a multifunctional detoxifying filter (MDF) capable of performing gas exchange and renal replacement in a single cartridge. Ex-vivo blood tests confirmed the ability of the MDF to oxygenate blood, remove carbon dioxide, and eliminate uremic toxins. In addition, animal experiments demonstrated the considerable clinical potential of this novel integrated extracorporeal life support approach. Integrating respiratory and renal support into a singular device could mitigate risks, conserve resources, and enhance the survival rates of critically ill patients suffering from concurrent lung and kidney failure.

---

Multi-organ failure is commonly encountered in ICUs, and recent studies have highlighted the interconnected nature of organ failures, particularly the effects of AKI on distant organs.^1,2^ Despite medical advancements, AKI continues to pose a substantial challenge, especially when coupled with ALI in critically ill patients. The coexistence of these conditions results in mortality rates exceeding 80%^3^, stressing the critical need for deeper understanding and targeted efforts to address the complex dynamics of MOF in the ICU environment.^4,5^

ECMO is being increasingly used to treat patients with refractory respiratory and/or circulatory failure. Single-organ dysfunction may be reversible given time for recovery. However, as the number and severity of organ failures increases, mortality rates also rise.^6^ Furthermore, a significant number of patients receiving ECMO also suffer from AKI and fluid overload, and in 50-60% of cases, CRRT is required.^7,8^ Accordingly, there is an urgent need to manage these organ failures more effectively. Several recent studies have explored the feasibility of integrating organ support therapies, including CRRT, ECMO, and extracorporeal CO_2_ removal (ECCO2R), ^9,10^ and the COVID-19 pandemic underscored the necessity for integrating renal replacement therapy (RRT) into ECMO circuits.^11^ Various approaches offer the possibility of ECMO and CRRT integration, such as separate vascular access and an RRT machine incorporating a hemofilter directly into the ECMO circuit using the blood flow provided by an ECMO (the “in-line technique”) or the utilization of two separate extracorporeal circuits (ECMO and CRRT) using an independent CRRT machine,^12^ both of which have shown to improve patient outcomes.^10,13^

However, despite the use of combined therapies, the overall survival rate remains disappointingly low at only 17%.^14^ Moreover, integrating CRRT and ECMO using these methods remains challenging as it necessitates separate filters and devices. This integration increases total artificial surface areas by introducing additional tubing and blood circuits, thus heightening the risk of platelet activation, adhesion, and aggregation and consequently increasing the risk of blood clot formation.^15^ Also, activation of the complement system and the release of inflammatory mediators can trigger sepsis, and the complexity of integrated blood circuits limits treatment to specialized medical centers. One study showed that the early initiation of CRRT increased the effectiveness of ECMO at avoiding fluid overload,^16^ demonstrating that acknowledgment of the ‘significance of effectively supporting multiple organ functions is essential for maximizing patient survival in the intricate context of ECMO and CRRT integration.^17^

Combining filters into a single cartridge and controlling the two therapies using one device can minimize patient/extracorporeal circuit interactions. Furthermore, an integrated device may be especially beneficial in spatially constrained and harsh environments where deploying separate devices might present logistical challenges. Also, this strategy enhances the utilization of available resources and paves the way for innovative solutions in scenarios such as those in air ambulances and during the COVID pandemic. Consequently, the Pneuma-K ECLS system features an MDF capable of performing two critical functions. Ex-vivo and animal experiments showed that this filter can supply oxygen and remove carbon dioxide from the bloodstream. In addition, the Pneuma-K ECLS system can support renal function by not only removing uremic molecules but also adjusting fluid balance.

## Materials and Methods

### The Pneuma-K ECLS system and the MDF

The Pneuma-K ECLS system consists of a Pneuma-K ECLS device and an MDF and was engineered to support impaired lung and kidney functions simultaneously.

The MDF, a core component of the Pneuma-K ECLS system, includes an inner housing for oxygenation and hemofiltration, a connector for housing placement, and caps to connect the housing and connector securely (**Fig. 1**). The hemofiltration and oxygenation filters are arranged coaxially within the core of the MDF and are linked by leak-proof connectors. The inner housings for the hemofiltration and oxygenation filters are cylindrical and incorporate semi-permeable membranes in the form of hollow tubes (fibers), which are potted in synthetic resin at both ends.

**Fig. 1.**
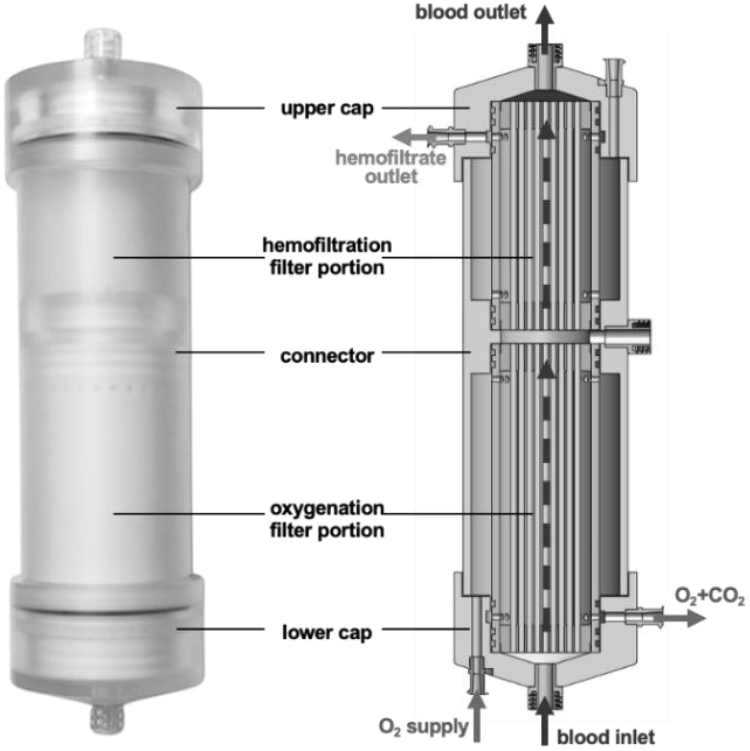
MDF (left) and internal flow diagram inside MDF (right)

The inner housings of the hemofiltration and oxygenation filters feature side holes in the circumferential direction. These side holes, which are positioned axially on both sides of the inner housing, facilitate fluid flow through the hemofiltration or oxygenation filter (**Fig. 1**).

Detailed specifications of the MDF are provided in **Table 1**. About 0.47 m^2^ of the membrane surface area is assigned to hemofiltration and 1.45 m^2^ of membrane surface area to the oxygenation filter.

**Table 1.**
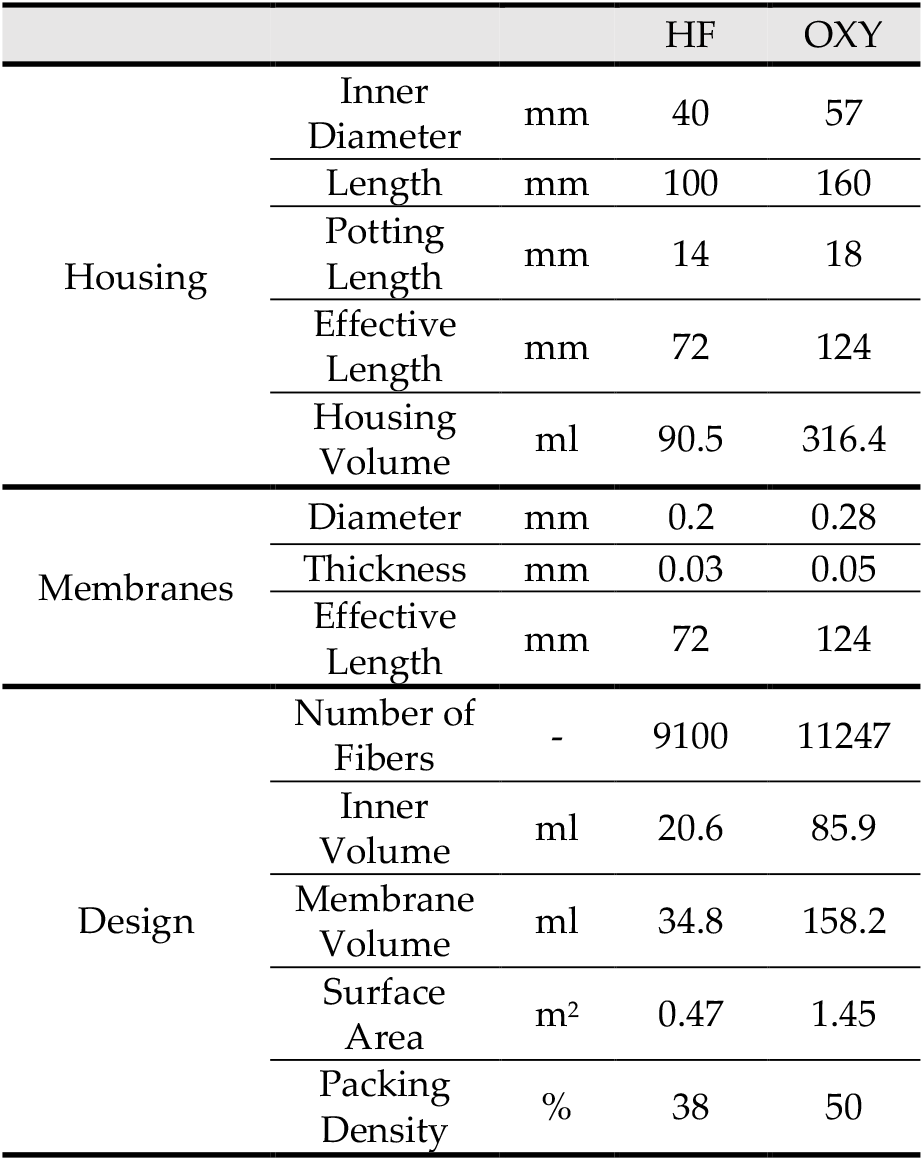
MDF Specifications (HF: hemofiltration, OXY: oxygenation)

|  |  |  | HF | OXY |
| --- | --- | --- | --- | --- |
| Housing | Inner Diameter | mm | 40 | 57 |
|  | Length | mm | 100 | 160 |
|  | Potting Length | mm | 14 | 18 |
|  | Effective Length | mm | 72 | 124 |
|  | Housing Volume | ml | 90.5 | 316.4 |
| Membranes | Diameter | mm | 0.2 | 0.28 |
|  | Thickness | mm | 0.03 | 0.05 |
|  | Effective Length | mm | 72 | 124 |
| Design | Number of Fibers | - | 9100 | 11247 |
|  | Inner Volume | ml | 20.6 | 85.9 |
|  | Membrane Volume | ml | 34.8 | 158.2 |
|  | Surface Area | m<sup>2</sup> | 0.47 | 1.45 |
|  | Packing Density | % | 38 | 50 |

The Pneuma-K ECLS device includes a B-pump to circulate blood, an S-pump for reinfusion, and a D-pump for hemofiltration. A heparin pump (H-Pump) is installed in the front of the device to enable the continuous infusion of anticoagulants upstream of the MDF. Pump flow rates are adjustable and can be calibrated using the display panel. The approximate dimensions of Pneuma-K ECLS device are 30 × 30 × 40 cm, not including the intravenous pole (**Fig. 2**).

**Fig. 2.**
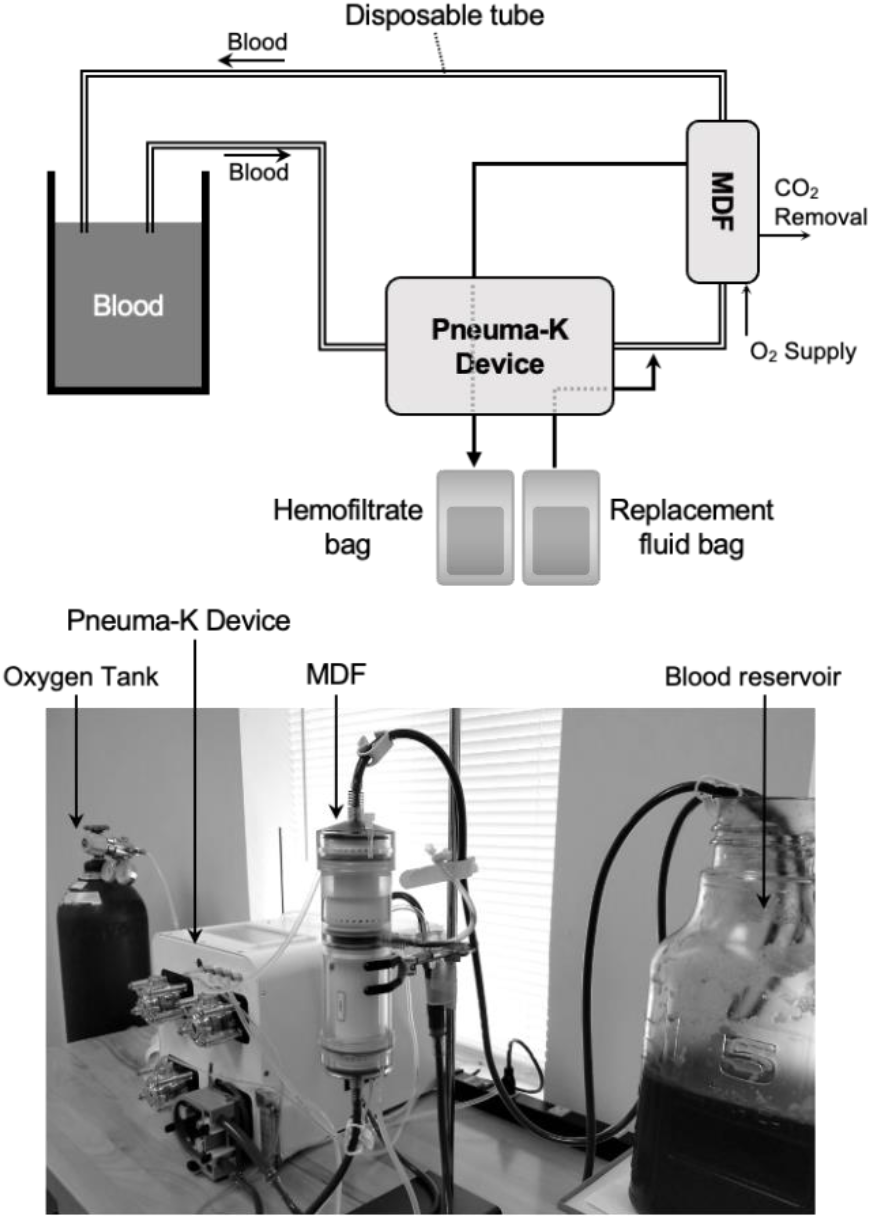
Ex-vivo circuit diagram

### Ex-vivo Blood Test Design

The Ex-vivo circuit is illustrated on **Fig. 2**. Fresh bovine blood (∼3 L), anticoagulated with heparin at 7.5-15 USP per ml of blood, was used as a blood substitute. Heparin was administered continuously or intermittently via the H-Pump to maintain anticoagulation throughout the experiments.

Oxygen was supplied at 700 ml/min at a pressure of 110 mmHg and was adjusted using a gas metering valve (Swagelok, USA) positioned downstream of the MDF. Blood flow was set at a maximum rate of 500 ml/min. Hemofiltration was performed at 30-35 ml/min, resulting in a total filtration volume of 10.8-12.6 L over 6 hours. To maintain fluid balance, an equivalent volume of replacement fluid (Prismasol BGK 2/3.5, Baxter) was reinfused into the bloodstream to maintain a net filtration rate of zero. The circuit was primed with isotonic saline (0.9% NaCl) heparinized at 5 USP per ml for 30 minutes before introducing blood.

To simulate kidney failure, uremic toxins (urea at 2.14 g/L and creatinine at 0.2 g/L) were dissolved in 100 ml of isotonic saline and added to the blood reservoir. In addition, fluorescein isothiocyanate-inulin (FITC-inulin, MW ∼5 kDa) was added at 0.2 g/L of blood. Hematocrit levels were maintained between 30-42%. During stabilization, an acute respiratory distress syndrome (ARDS) model was prepared by preconditioning blood substitute with CO_2_. A CO_2_ tank was connected to the outer oxygenation membranes of the MDF to deliver CO_2_; partial CO_2_ pressure was increased to over 130 mmHg.

CO_2_ clearance through the MDF was quantified under various pre-MDF pCO_2_ conditions, ranging from 50 to 90 mmHg. Blood samples were collected from the blood reservoir and upstream and downstream of the MDF at t=0, 1, 2, 4, and 6 hours. About 10 ml of blood was withdrawn from each site for blood gas analysis; a VetScan i-STAT (Abaxis, USA) with a CG8+ Cartridge was used to analyze Na+, K+, Ca++, Glucose (Glu), Hematocrit (Hct), Hemoglobin (Hgb), pO_2_, sO_2_, pH, pCO_2_, HCO_3_, TCO_2_, and Base Excess levels.

### Animal Experiment Protocol

The Institutional Animal Use and Care Committee (IACUC) at the University of Vermont approved the animal protocol and all studies were performed under institutional and Association for Assessment and Accreditation of Laboratory Animal Care (AAALAC) guidelines. Six adult female pigs (40-50 kg) were anesthetized with ketamine (20-30 mg/kg) and atropine (0.05-0.5 mg/kg) and then continuously administered isoflurane (1-5%). The experiment involved inducing partial lung dysfunction (hypercapnic respiratory failure) by adjusting FiO_2_ to 21% and reducing the respiratory rate to 4.5±0.5 breaths/min, aiming for SaO_2_ ≤ 60% and pCO_2_ ≥ 58 mmHg.

A 16 French triple-lumen catheter was inserted into the external jugular vein for central venous access. Catheters were positioned above the heart in the superior vena cava, and their placements were confirmed by fluoroscopy. Heparin was administered at 300 USP/kg, and activated clotting time (ACT) was monitored about 10 minutes later. A 150 USP/kg heparin bolus was administered when ACT fell below 350 sec, and ACT was measured every 10 minutes until it exceeded 350 sec.

The Pneuma-K ECLS system was initiated when target pCO_2_, SaO_2_, and ACT levels were reached. After system initiation, heparin was infused continuously at an initial rate of 20 USP/min. ACT levels were monitored and maintained between 200 and 250 sec, and when the level fell below 200 sec, a 150 USP/kg heparin bolus was administered, or the continuous heparin infusion was increased to 100 USP/min.

The Pneuma-K ECLS system provided a blood flow rate of 500 ml per minute with a sweep gas flow of 5 or 10 L per minute. Blood pressure and oxygen saturation were continuously monitored by ear and femoral artery cannulation. Blood samples (10 ml) were taken hourly from the venous cannula and analyzed for CBC, electrolytes, renal function, and coagulation. In addition, blood was withdrawn from the arterial cannulas for arterial blood gas analyses (pH, pO_2_, pCO_2_, HCO_3_-, and O_2_ saturation).

## Results

### Ex-vivo Blood Tests

Six blood tests were conducted using the MDF system, and no significant malfunctions were observed. CO_2_ clearance was effective, with pCO_2_ levels decreasing from 86.7 to 9.4 mmHg in the blood reservoir over 6 hours (**Fig. 3**). Under controlled upstream conditions, pCO_2_ levels were reduced significantly through the MDF; pre-MDF pCO_2_ levels of 85.3±2.8, 65.6±2.5, and 57.1±2.1 mmHg were reduced to post-MDF levels of 62.8±4.5, 48.2±5.3, and 44.9±4.1 mmHg, respectively. In parallel, pO_2_ levels in the blood reservoir progressively increased to over 300 mmHg a few minutes after introducing oxygen, and oxygen saturation levels also increased within minutes (**Fig. 3**). These results confirmed that the MDF provided effective gas transfer.

**Fig. 3.**
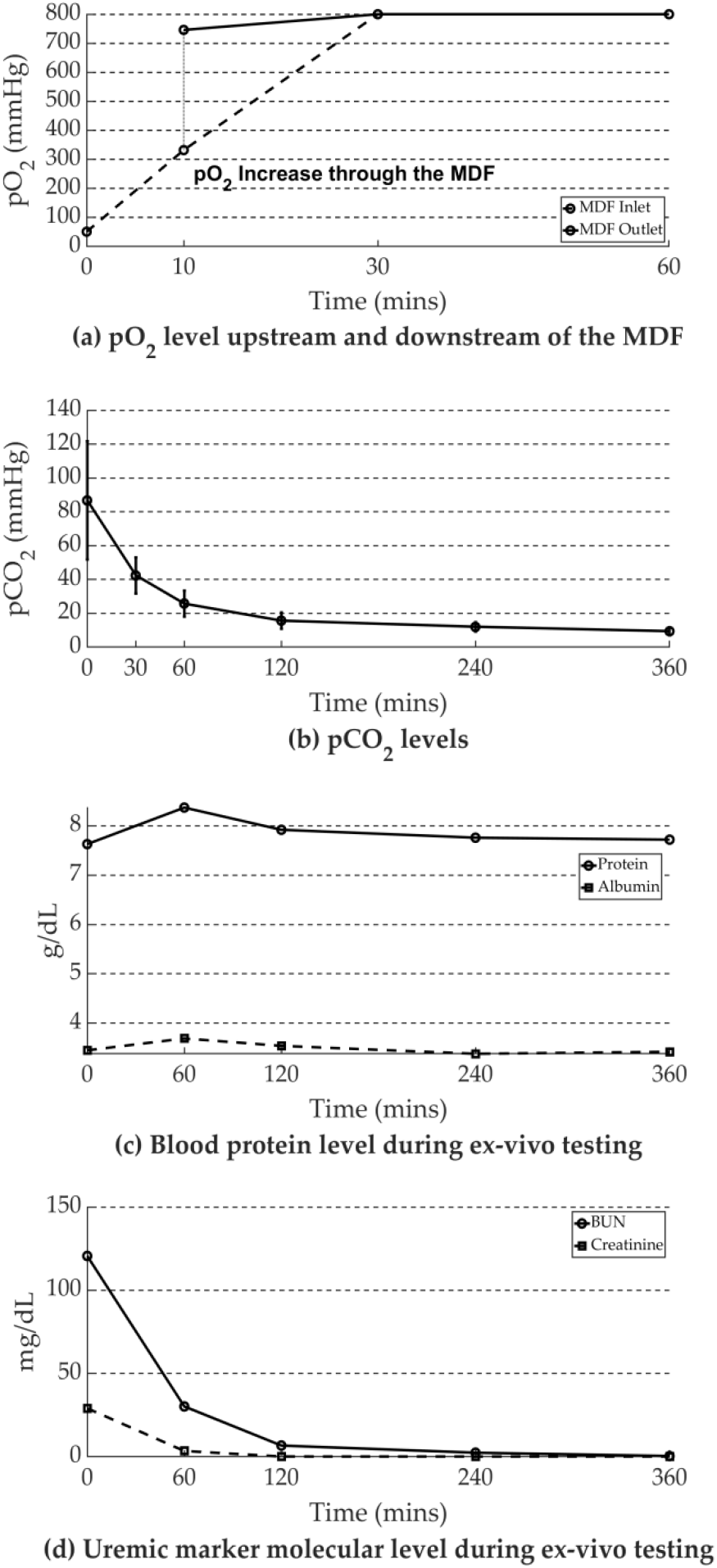
Ex-vivo Blood Test Results: (a) pO_2_ level upstream and downstream of the MDF; (b) pCO_2_ levels; (c) Blood protein levels during Ex-vivo testing; (d) Uremic marker molecular level during Ex-vivo testing

In the simulated renal failure model, uremic molecule levels were efficiently reduced with the MDF system; a 99% reduction in urea levels was obtained over the 6-hour test period. Creatinine levels showed a similar reduction; no significant protein or albumin loss occurred (**Fig. 3**).

### Animal Experiments

Anticoagulation was adequately managed during the animal experiment, and anesthesia was effectively maintained. Moreover, there were no signs of MDF leakage or malfunction (**Fig. 4**). The Pneuma-K ECLS system supported subjects with partial lung dysfunction for an average of 9±1.5 hours during experiments that lasted up to 12.5 hours. No notable difference in hematocrit levels or any increase in plasma free hemoglobin (pHb) levels was observed after 12 hours of Pneuma-K ECLS operation.

**Fig. 4.**
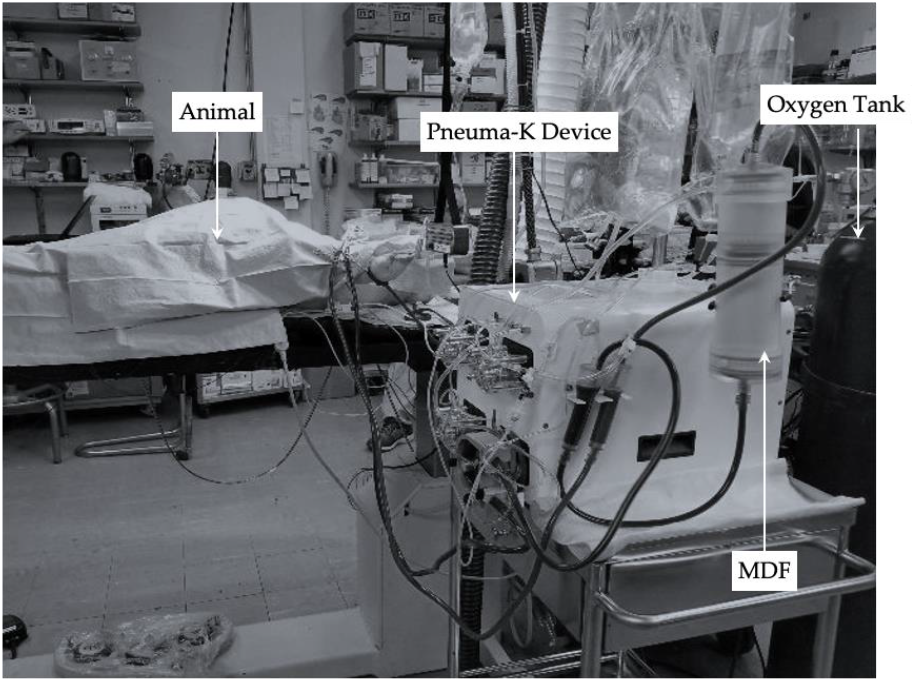
Animal Experiment

**Table 2** summarizes pre- and post-MDF blood gas analysis results from three animal experiments. SaO_2_ reached 100%. The MDF substantially increased pO_2_ levels, and pCO_2_ levels were reduced from 58.1±3.3 to 35.9±4.2 mmHg. Marked differences in blood colors upstream and downstream of the MDF confirmed adequate oxygenation and CO_2_ removal. During treatments, renal function markers, including creatinine levels, remained stable, and total protein levels were maintained without significant loss. (**Fig. 5**)

**Table 2.** Blood gas analysis results upstream and downstream of the MDF during animal tests.

|  | Pre MDF | Post MDF |
| --- | --- | --- |
| $pCO_2$ (mmHg) | $58.1 \pm 3.3$ | $35.9 \pm 4.2$ |
| $pO_2$ (mmHg) | $39.7 \pm 3.8$ | $366.7 \pm 142.9$ |
| $SaO_2$ (%) | $68.7 \pm 8.3$ | $100.0 \pm 0$ |
| $HCO_3$ (mmol/L) | $31.9 \pm 1.3$ | $30.8 \pm 0.5$ |
| pH | $7.3 \pm 0.1$ | $7.50 \pm 0.1$ |

**Fig. 5.**
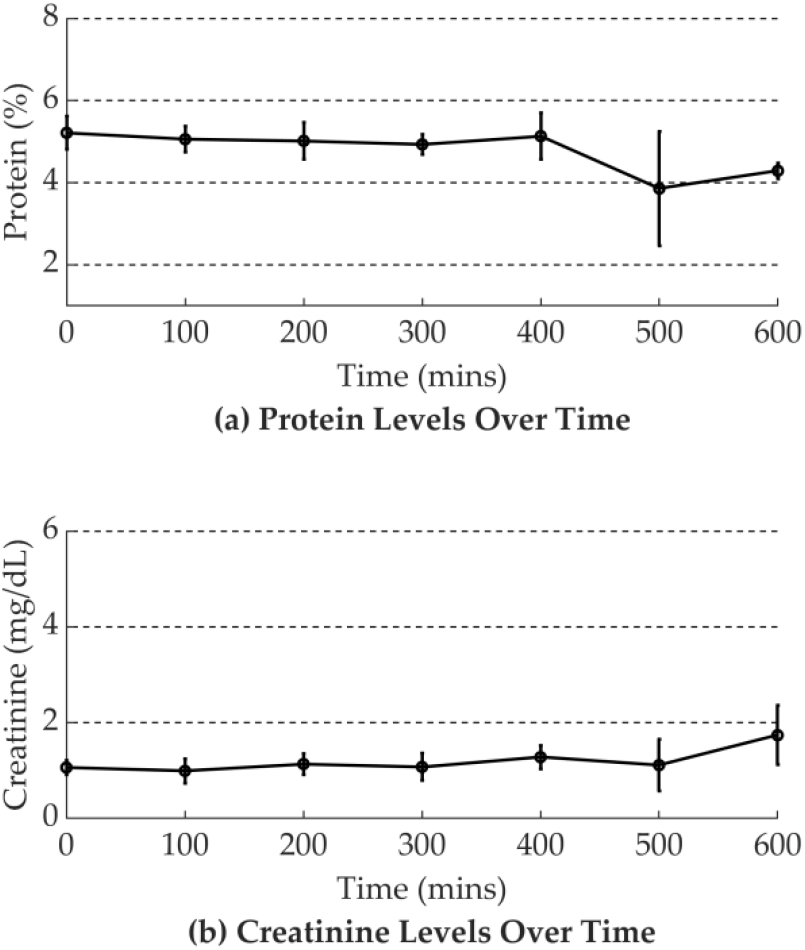
Longitudinal Animal Experiment Results: (a) Protein levels; (b) Creatinine levels

## Discussion

This study investigated the feasibility of combining two separate extracorporeal organ support systems into a single unit. The design of this device and filter constitutes a significant technological advancement and might enhance the treatment of MOF patients considerably by improving treatment efficiency and convenience in critical care settings. In addition, integrating functions reduced the number of components, which decreased blood flow resistance and minimized the risks of contamination and coagulation while enabling more streamlined operation.

Extensive Ex-vivo and animal experiments demonstrated that the MDF effectively removes CO_2_ while oxygenating blood, as confirmed by pCO_2_, pO_2_, and sO_2_ levels. Enhanced blood oxygenation was also evidenced by visible changes in blood color upstream and downstream of the MDF. In addition, tests demonstrated that the MDF positively affected renal support.

Although this study demonstrates the clinical advantages of the MDF, there is scope for further optimization to enhance the effectiveness and safety of the device. For instance, altering the surface area of the hemofiltration and oxygenation filter components could improve the efficiency of gas exchange (oxygenation) and mass transfer (hemofiltration). Currently, the Pneuma-K ECLS system includes 1.45 m^2^ for oxygenation and 0.47 m^2^ for hemofiltration, whereas commercial filters provide 2 to 4 m^2^ of effective surface area for ECMO and 0.6 to 2.5 m^2^ for CRRT. In addition, gas transfer in the current system is slower than in conventional ECMO setups. Increasing the effective surface area of the membrane dedicated to oxygenation might address this issue.

Fiber arrangements might also provide a means of improving efficiency. An “outside-in” fiber arrangement is employed in typical ECMO systems, wherein blood flows through the outer hollow fiber compartment.^18^ However, the current oxygenation part of the MDF utilizes an “inside-out” arrangement, that is, blood flows through the interiors of the hollow fibers, and the “outside-in” configuration may be superior in terms of the gas exchange.^19^ Moreover, the “outside-in” approach could address issues of fiber clogging and the shorter filter lifespans of “inside-out” filters.^26^ Consequently, redesigning the MDF to facilitate blood flow outside the fiber membranes might prove beneficial.

The Pneuma-K ECLS system allowed a blood flow rate of 500 ml/min to be maintained throughout experiments. CRRT typically necessitates a blood flow of 100 to 200 ml/min,^20^ whereas ECMO requires a much higher rate of 2 L/min ^21^. The lower rate required for CRRT is selected for patients who may be hemodynamically unstable and provides a more gradual, gentle extraction of waste products and excess fluid from the bloodstream.^22^ When integrating CRRT and ECMO into a single MDF unit, a compromise flow rate of 500 ml/min was selected to balance hemodynamic stability and gas/mass transfer efficiency. This rate fell within the optimal range of 400 to 500 ml/min suggested for CRRT and extracorporeal CO_2_ removal.^22^ In addition, a rate of 500 ml/min rate was deemed suitable because it falls within the range used for pediatric ECMO, which varies from 200 ml to 2.8 L.^23^ However, the 500 ml/min blood flow rate resulted in slower gas transfer than traditional ECMO devices, indicating the need for further MDF optimization. Further study is required on the effects of higher blood flow rates on system and filter capacity.

Filter design is versatile and can accommodate diverse membrane combinations, making it suitable for treating many MOF types, including liver failure and sepsis. Individual organ dysfunctions can be specifically targeted, and the Pneuma-K ECLS device can be customized according to individual patient needs by tailoring membrane configurations.^24^ This adaptability not only extends the applicability of the device beyond initial design intentions but also enhances its effectiveness as a solution for various organ support requirements. Ultimately, this flexibility could profoundly impact the field of critical care by improving both patient care and outcomes.^24^

Finally, the lifespan of the filter needs consideration. We conducted animal experiments spanning approximately 12 hours, yet ECMO patients frequently require extended treatment durations, often exceeding several days ^25^. Therefore, it is essential to pursue further research on the long-term use of the MDF to determine its operational lifespan.

## Conclusion

The Pneuma-K ECLS system simplifies caregiver workflow, minimizes required training, and enables the standardization of protocols. These advantages facilitate system optimization, reduce complexities, and improve the efficiency and accessibility of multi-organ support to enhance patient care and outcomes.

## Notes

### Competing Interest Statement

The authors have declared no competing interest.

